## Supplemental Figures 1-13 for "Joint profiling of DNA and proteins in single cells to dissect genotype-phenotype associations in leukemia"

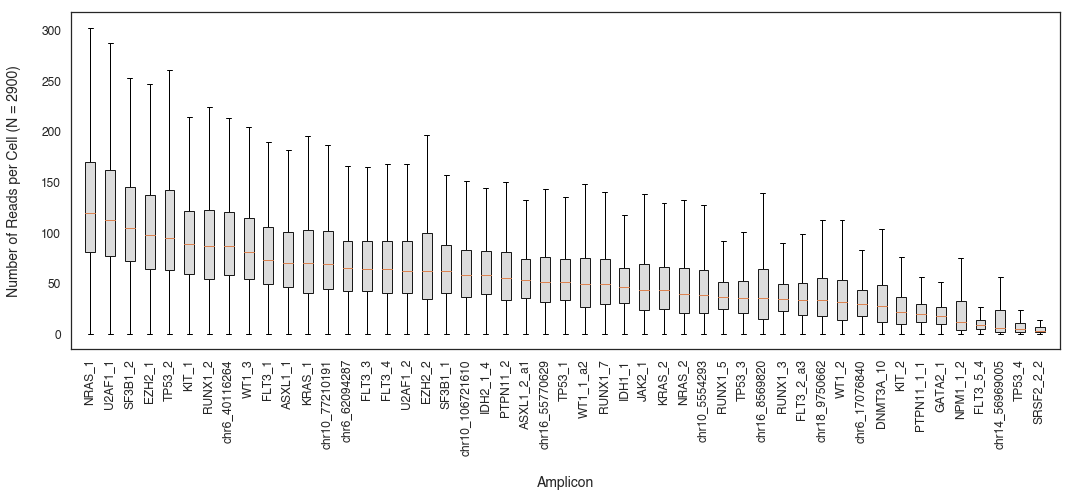

**Supplementary Figure 1: Reads per DNA target for the 3-cell control experiment.**

Distribution of reads per cell for each of the 49 DNA panel targets are presented as a boxplot and sorted in descending order by median number of reads per cell (orange bar).

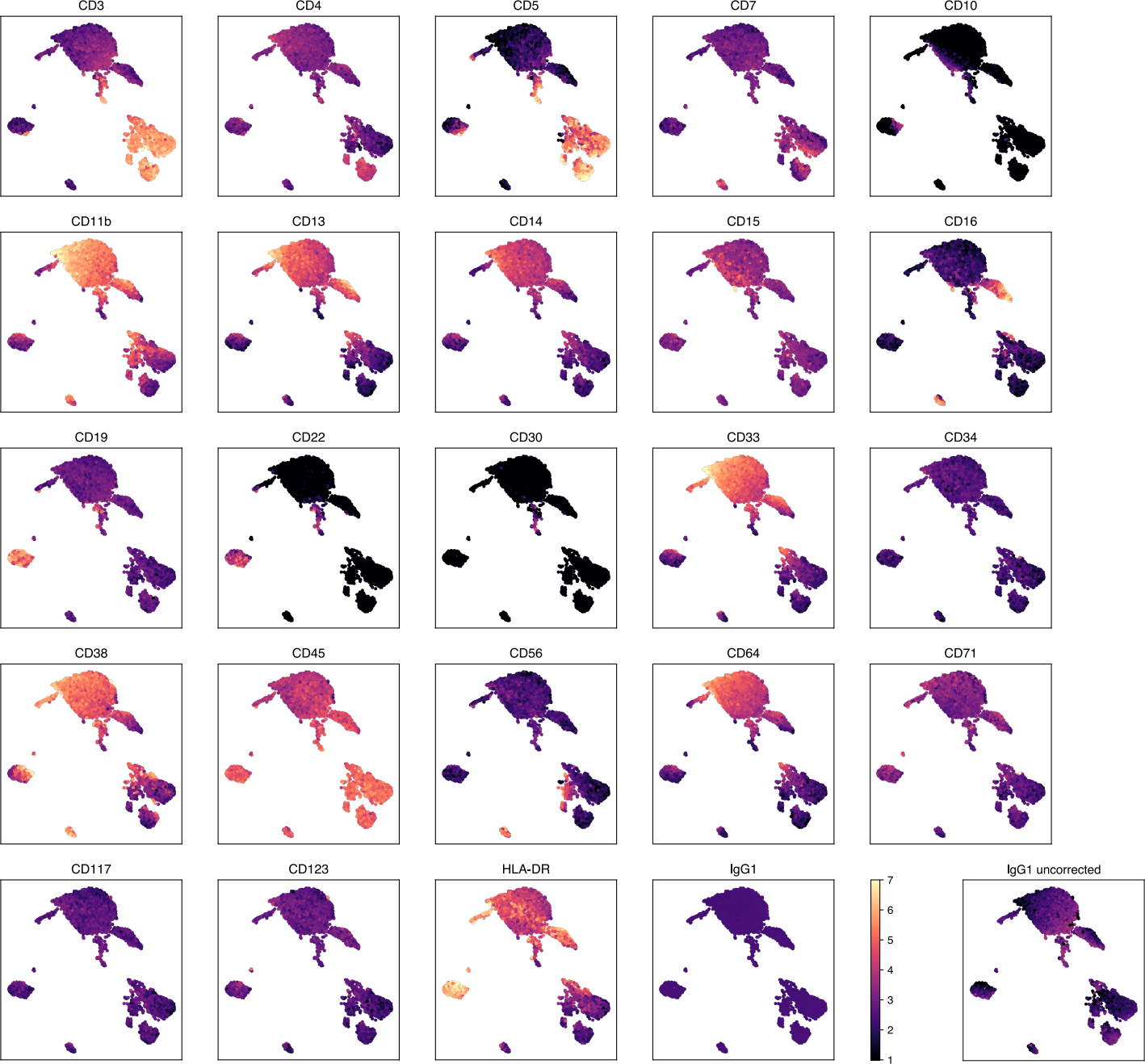

**Supplementary Figure 2: Antibody signal distribution in the PBMC control.**

Corrected antibody counts (log scale with base *e*) for each cell and antibody are given as a heatmap using the UMAP coordinates from Figure 2. Heatmap values are clipped at 1 and 7. Since IgG1 signal was used in the antibody correction as a co-regressor together with other size factors (see Methods), the residual count values are uniform. Therefore, the non-regressed log-transformed IgG1 counts are provided in an additional panel (lower-right).

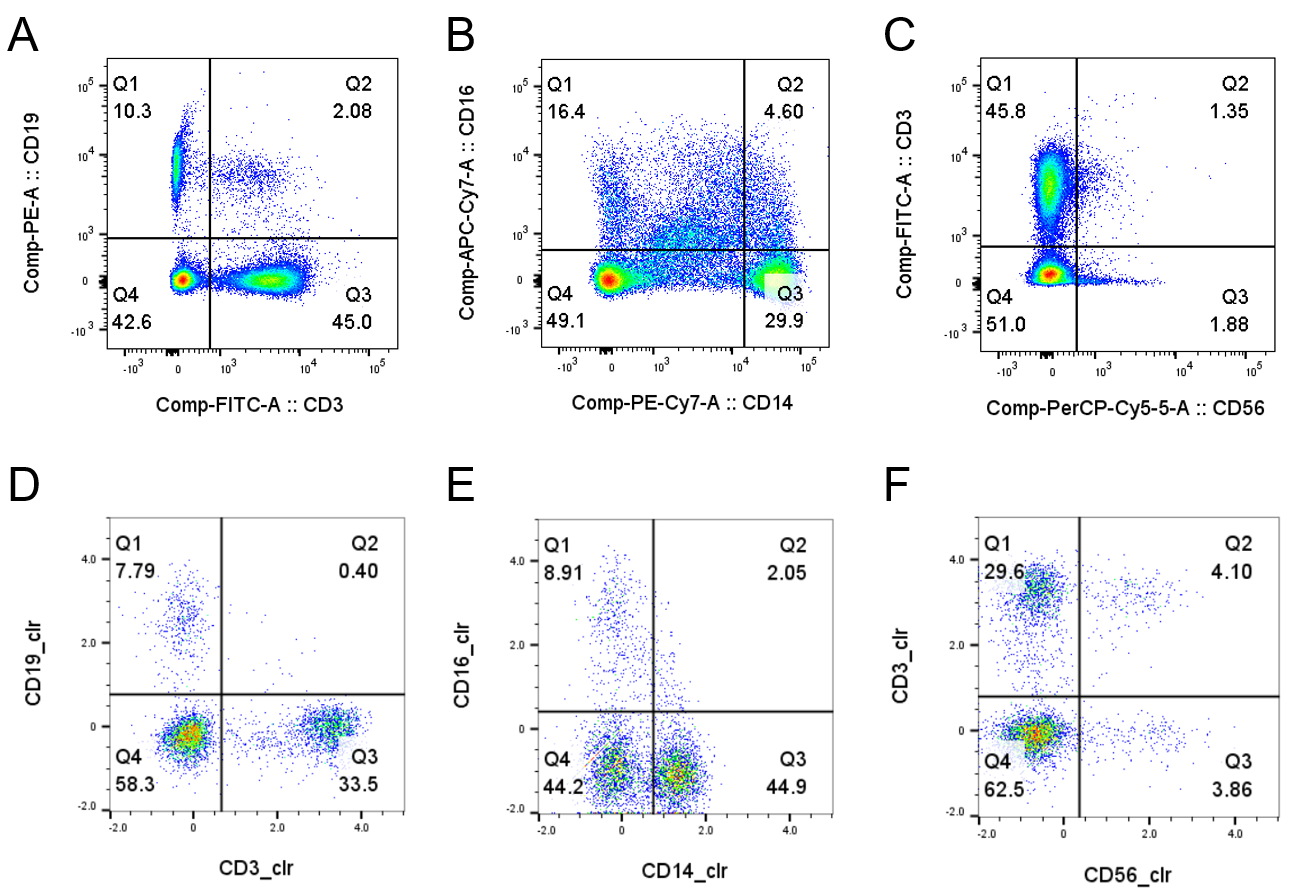

**Supplementary Figure 3: Comparison of DAb-seq data to flow cytometry for healthy PBMCs.**

**(A, B, C)** Flow cytometry results using five hematopoietic markers (CD3, CD14, CD16, CD19, CD56) to discriminate blood cell populations in healthy PBMCs. **(D, E, F)** DAb-seq results for the same PBMC sample across the five markers. Antibody signal is expressed as the centered log ratio for each cell.

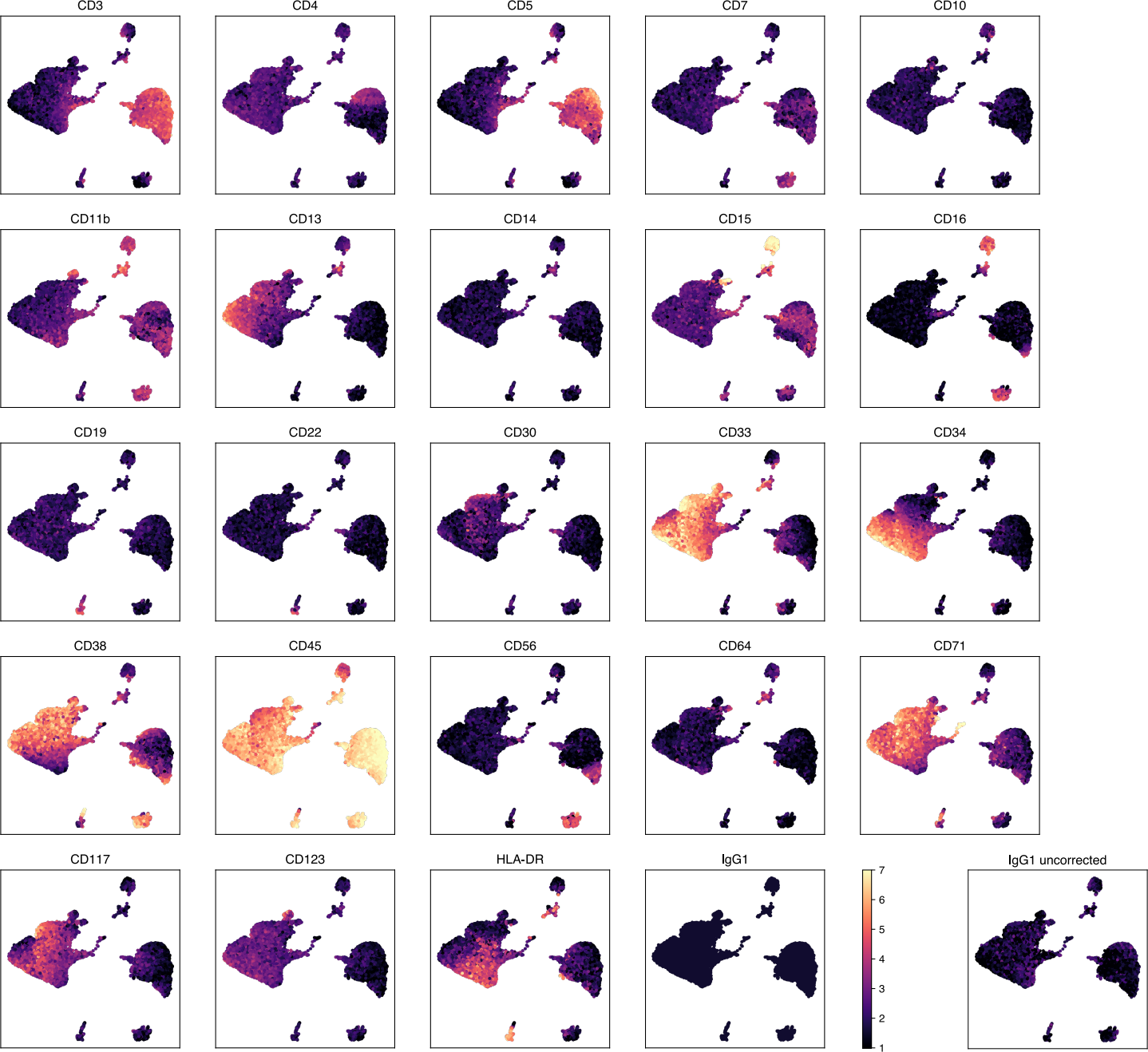

**Supplementary Figure 4: Antibody signal UMAP for Patient #1 (gemtuzumab treated).**

Corrected antibody counts (log scale with base *e*) for each cell and antibody are given as a heatmap using the UMAP coordinates from Figure 3. Non-regressed log-transformed IgG1 counts are provided in an additional panel (lower-right).

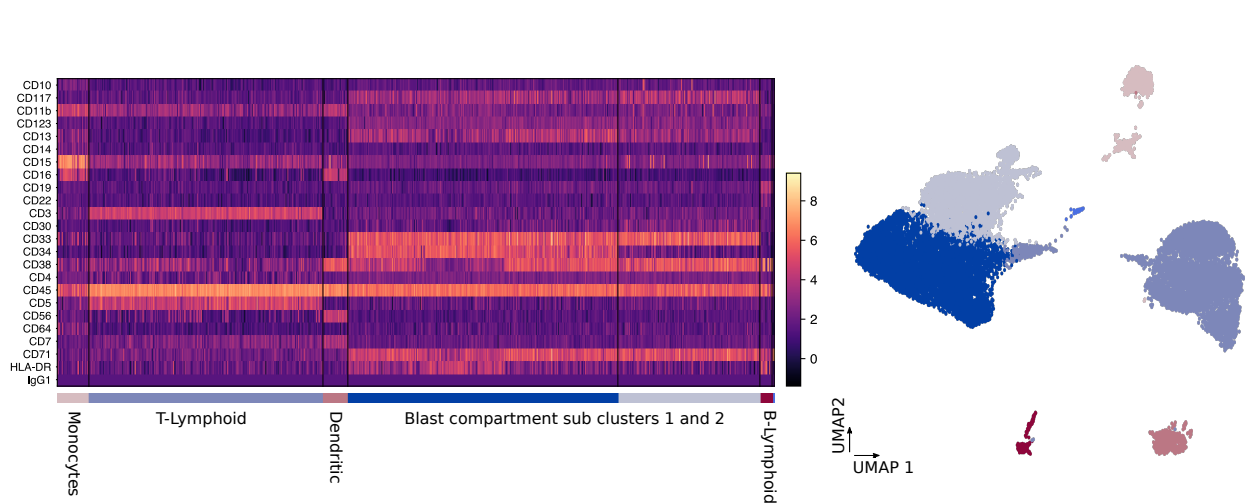

**Supplementary Figure 5: Antibody count heatmap for Patient #1 (gemtuzumab treated).**

Corrected antibody counts (log scale with base *e*) are represented as a heatmap and sorted by the cell compartment clusters as described in the main text (Leiden clustering of the corrected antibody count matrix at a resolution factor of 0.4).

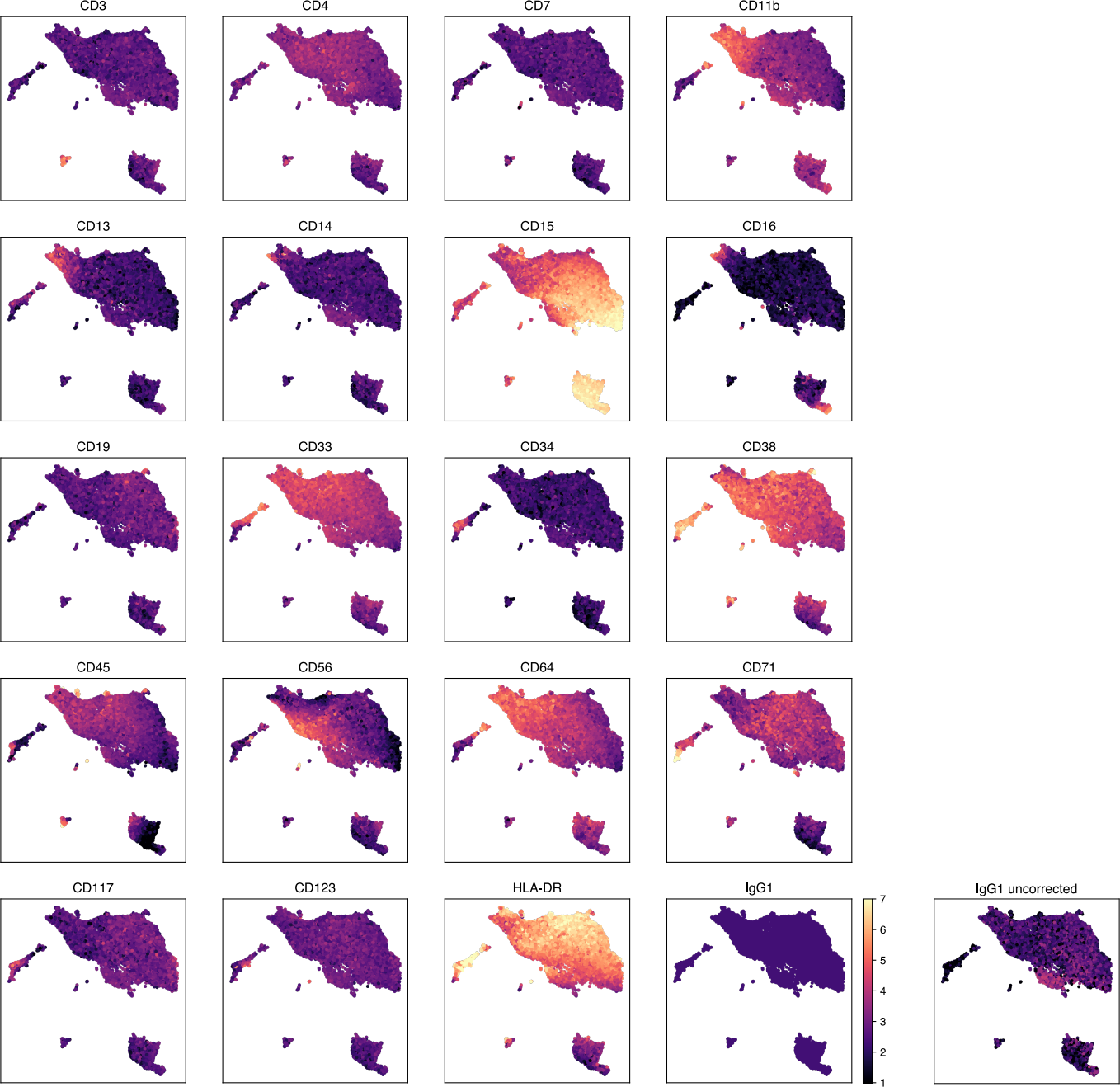

**Supplementary Figure 6: Antibody signal UMAP for Patient #2 (pediatric patient).**

Corrected antibody counts (log scale with base *e*) for each cell and antibody are given as a heatmap using the UMAP coordinates from Figure 4. Non-regressed log-transformed IgG1 counts are provided in an additional panel (lower-right).

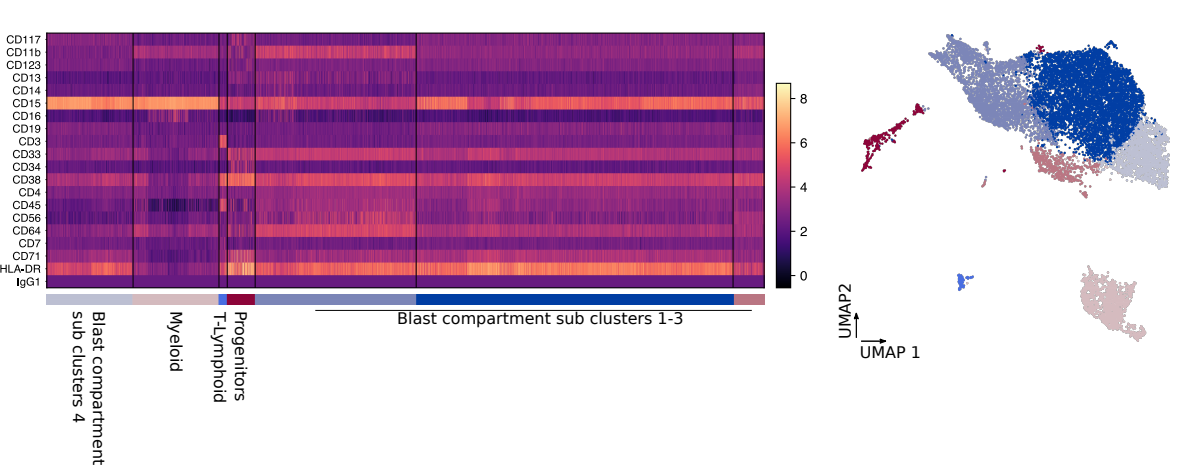

**Supplementary Figure 7: Antibody count heatmap for Patient #2 (pediatric patient).**

Corrected antibody counts (log scale with base *e*) are represented as a heatmap and sorted by the cell compartment clusters as described in the main text (Leiden clustering of the corrected antibody count matrix at a resolution factor of 0.4). Blast subclusters 1-3 are not well separated.

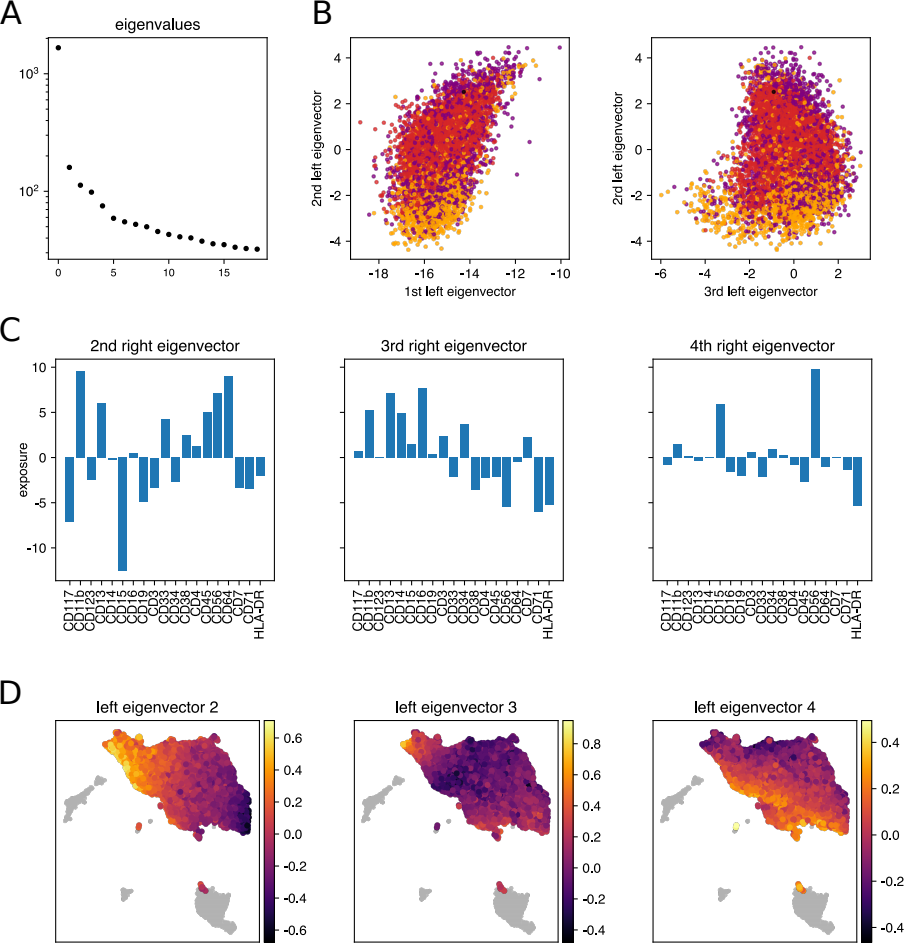

**Supplementary Figure 8: Singular value decomposition of blast cell compartment for Patient #2 (pediatric patient).**

Continuous trends are not easily identified by clustering. To discover such gradients in the antibody expression of the blast cells, singular value decomposition was calculated for the log-transformed and corrected cell by antibody counts matrix (**A = U * s * V^T^**) (blast cells only. No mean centering or additional normalization was performed). **(A)** Ordered singular values are given. **(B)** Scatter plot of individual cells in coordinates of the first three left singular vectors (scaled by the corresponding singular value, that is: **U_i_ * s_i_**). Colors are according to the genotype, same as in figure 4. **(C)** Second to fourth right singular vectors are given (again scaled by the corresponding singular value: **s_i_ * V^T^_i_**) which show the contribution of each antibody to the principal component. For example, CD15 and CD117 are anticorrelated with the second principal component (2^nd^ left eigenvector), and CD11b, CD13, CD56 and CD64 are correlated. The first right eigenvector is not plotted. Because antibody counts are all positive numbers, the first principal component of the decomposition corresponds to the count offset and therefore carries no useful gradient information. **(D)** Scatter plot of cells in same UMAP coordinates as in figure 4. Each cell within the blast compartment is colored according to its 2^nd^, 3^rd^, or 4^th^ left eigenvector value and highlights antibody count gradients within the blast compartment.

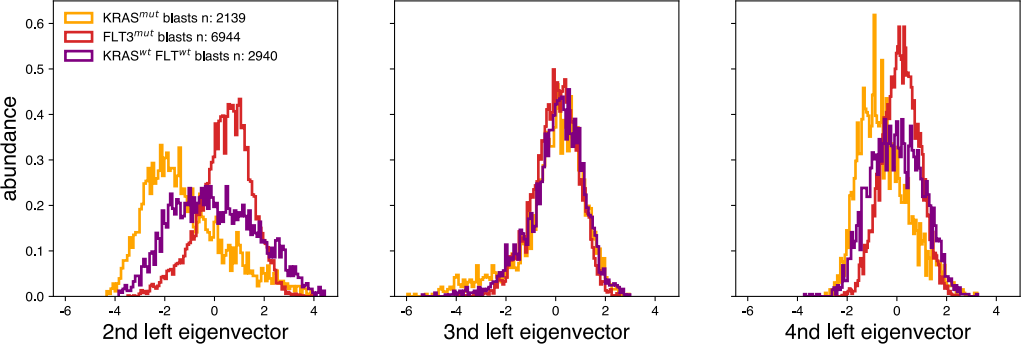

**Supplementary Figure 9: Blast genotype distribution along the 2^nd^, 3^rd^, and 4^th^ left eigenvectors for Patient #2 (pediatric patient).**

Distribution of cells within the blast compartment along the 2^nd^, 3^rd^, and 4^th^ left eigenvectors are shown. Histograms are plotted for each detected genotype separately. This shows that the genotype distribution within the blast compartments are biased toward a phenotypic cluster, but do not separate completely. In consequence, no FACS gating strategy can successfully deliver a genetically pure cell sample.

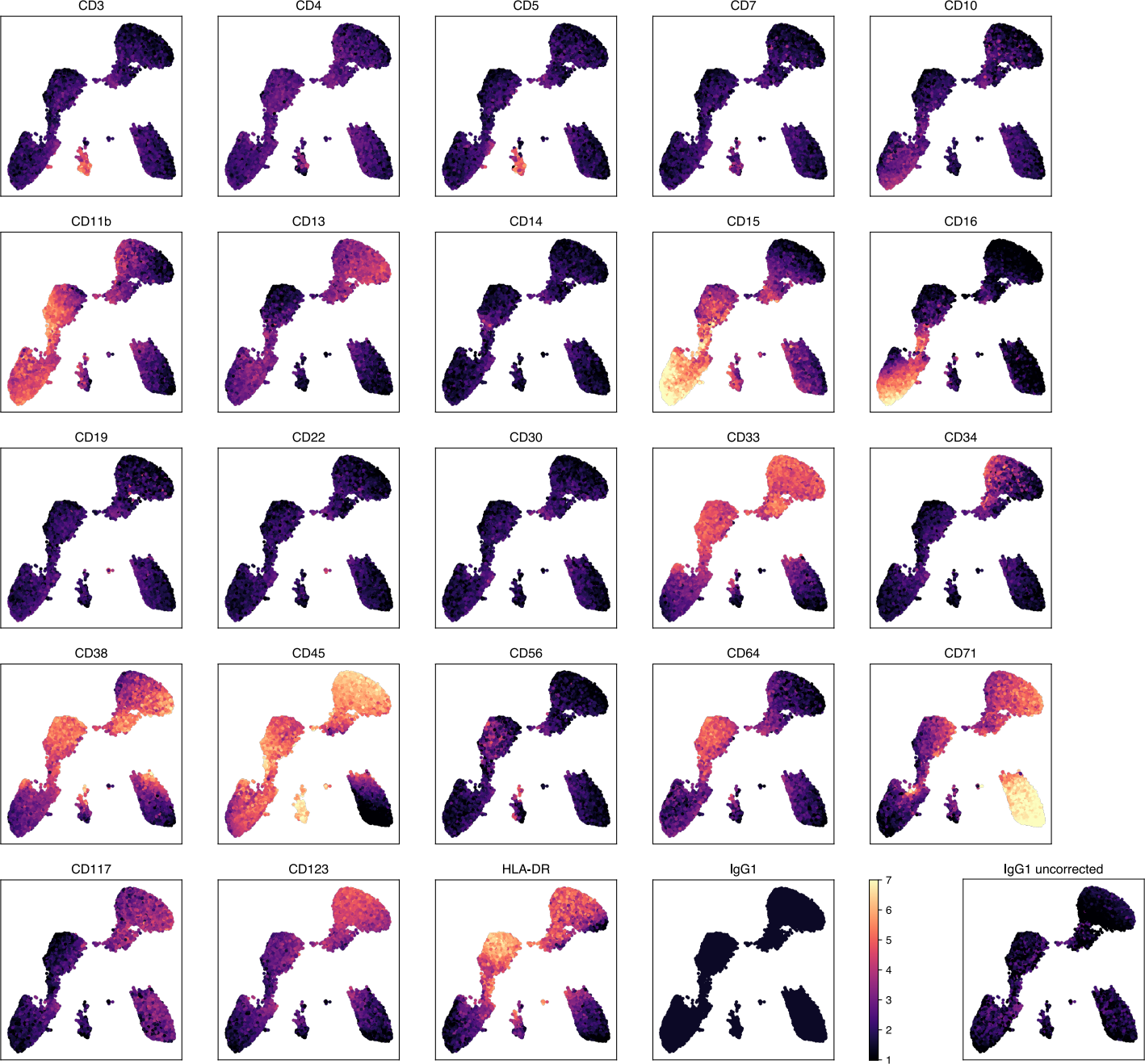

**Supplementary Figure 10: Antibody signal UMAP for Patient #3 (FLT3 inhibitor treated).**

Corrected antibody counts (log scale with base *e*) for each cell and antibody are given as a heatmap using the UMAP coordinates from Figure 5. Non-regressed log-transformed IgG1 counts are provided in an additional panel (lower-right).

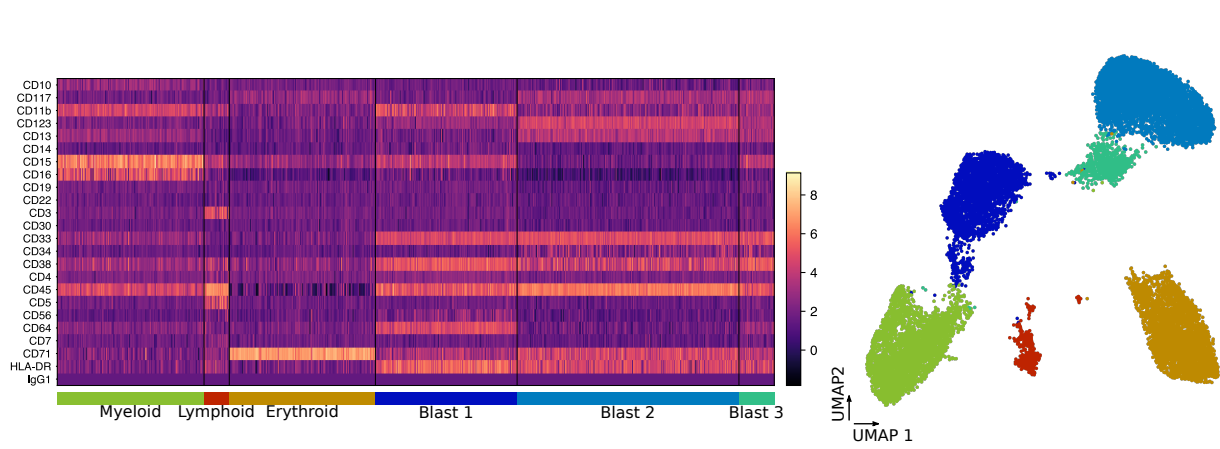

**Supplementary Figure 11: Antibody count heatmap for Patient #3 (FLT3 inhibitor treated).**

Corrected antibody counts (log scale with base *e*) are represented as a heatmap and sorted by the cell compartment clusters as described in the main text (Leiden clustering of the corrected antibody count matrix at a resolution factor of 0.4). Blast subclusters 1-3 are not well separated. Different lymphoid compartments are not labeled in the heatmap due to small cell numbers (e.g. B-lymphocytes and NK-lymphocytes), however, the clusters are visible in the UMAP representation.

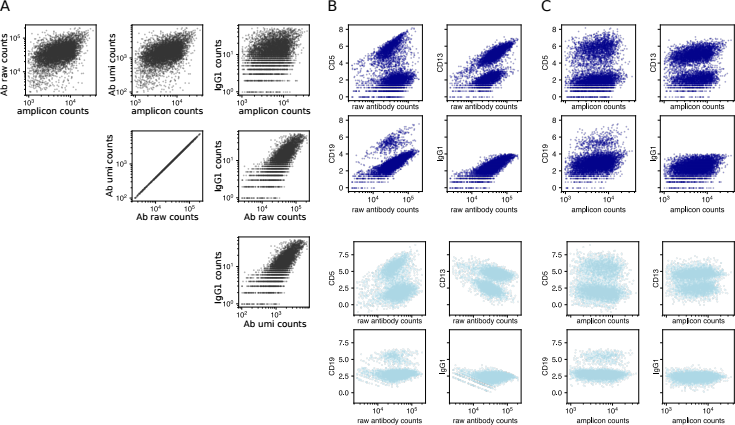

**Supplementary Figure 12: Antibody count correction performance.**

**(A)** Correlation among different cellular quality metrics (total antibody raw count, total antibody UMI count, amplicon count, total IgG1 count) are given as scatter plot for all cells in the PBMC control experiment. **(B)** For each cell in the PBMC experiment, raw antibody count (dark blue, top panel) or corrected antibody count (light blue, bottom panel) for four markers is plotted against the total raw antibody count per cell. Correlation in raw counts with total antibody count is clearly visible and present in the isotype control. Linear regression with the quality metrics as covariates reduces this dependency (light blue, bottom panel). **(C)** Same as (B) but plotted against total DNA panel read count. The correlation between total DNA panel count and individual antibody counts is weaker but nevertheless present.

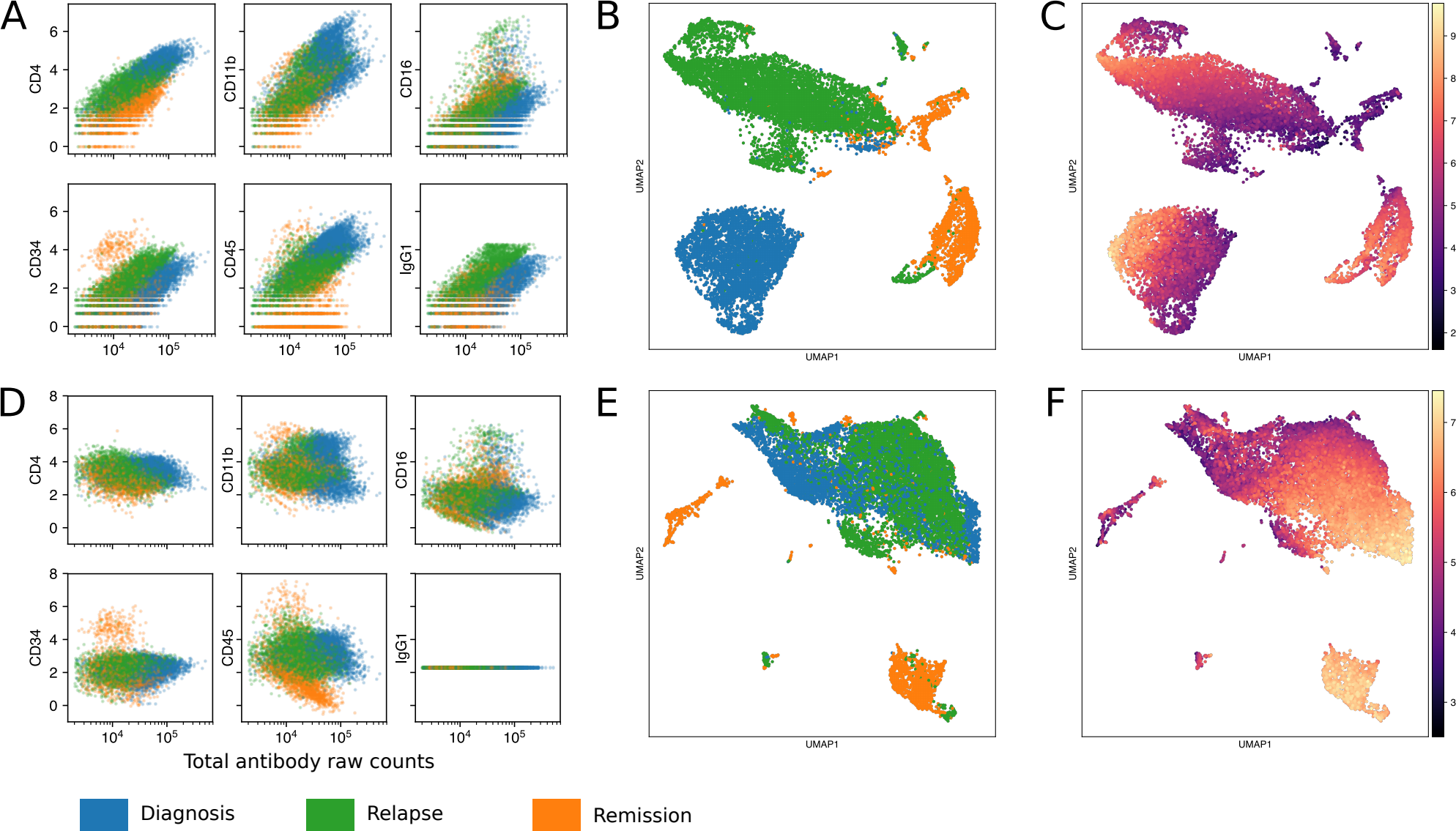

**Supplementary Figure 13: Simultaneous antibody count regression corrects for batch effects.**

**(A)** Six representative antibodies are plotted versus uncorrected total antibody counts for each cell in the Patient #2 (pediatric patient) series. Colors indicate sample timepoint. Batch effects are visible as an offset in the y-axis. **(B)** UMAP embedding of the uncorrected log transformed antibody counts is given. Color indicates sample timepoint. **(C)** Same UMAP embedding as in (B) but colored according to log(raw CD15 count). **(D)** Same as (A) after antibody count correction. Batch effect is reduced compared to (A). **(E)** Same as (B) but using corrected antibody counts to construct the UMAP embedding. After antibody correction, blast cells cluster together. **(F)** Same as (C) for corrected antibody counts. (E) and (F) correspond to the UMAP depicted in Figure 4.
